# Role of Anti-SARS-CoV-2 antibodies in different cohorts: Can they provide clues for appropriate patient triaging?

**DOI:** 10.1101/2020.06.26.174672

**Authors:** Manohar B. Mutnal, Amin A. Mohammad, Alejandro C. Arroliga, Yinan Hua, Liping Wang, William Koss, Arundhati Rao

**Author notes:** Address for correspondence, Manohar B. Mutnal, Ph.D., Section Chief – Microbiology, Department of Pathology and Laboratory Medicine, 2401, South 31^st^ street, Temple TX 76508.

## Abstract

The emergence of coronavirus disease 2019 (COVID-19) has become a major global health crisis. Currently, diagnosis is based on molecular techniques, which detect the viral nucleic acids when present at detectable levels. The serum IgG response against *SARS-CoV-2* was examined by using an ELISA-based assay. Serum samples, along with nasopharyngeal specimens were collected from various cohorts and analyzed by ELISA and rRT-PCR, respectively. A total of 167 serum samples were tested for serum IgG antibodies against *SARS-CoV-2* in outpatient cohorts, 15 (8.9%) were positive by rRT-PCR and the remaining 152 (91%) were negative. We used these data to generate two different assay cutoffs for serum IgG assay and investigated percent concordance with rRT-PCR test results. The emergency department data revealed, out of 151 nasopharyngeal swabs, 4 (2.6%) were positive by rRT-PCR and 18 (11.9%) were positive for serum IgG assay. Among the 18 patients that were positive for serum IgG, 13 (72.2%) exhibited 1-3 symptoms of COVID-19 and 5 (27.7%) patients did not present with any COVID-19 related symptoms, per CDC criteria. All 4 (100%) patients that were positive by rRT-PCR had symptoms of COVID-19 disease. A longitudinal study from the inpatient population suggested there was a sharp increase in the serum IgG titers in 5 patients, a moderate increase in 1 patient and a plateau in 3 patients. Sero-prevalence of COVID-19 disease in pre-procedure patients was 5.5%. Our findings suggest serological tests can be used for appropriate patient triaging when performed as an adjunct to existing molecular testing.

## Introduction

In December 2019, a series of pneumonia cases of unknown cause emerged in Wuhan, Hubei, China, with clinical presentations greatly resembling viral pneumonia^1^. Subsequently, pathogenic gene sequencing identified the infecting pathogen as a novel coronavirus, named Severe Acute Respiratory Syndrome Coronavirus-2 *(SARS-CoV-2)*^2^. It has been listed as a public health emergency of international concern, and, following declaration of a pandemic by the World Health Organization (WHO), governments worldwide have taken drastic measures to contain the outbreak, including the quarantine of millions of residents in many countries.

According to the Centers for Diseases Control and Prevention (CDC, Atlanta, GA, USA) and recent reports, most COVID-19 patients have an incubation period of 2 to 14 days^3^. CDC has listed 11 symptoms for clinical diagnosis. Fever, cough, shortness of breath and fatigue are the most common symptoms, whereas nasal congestion and diarrhea are only noted in a small number of patients^4^ Severe cases might progress to acute respiratory distress syndrome (ARDS), septic shock and difficult-to-tackle metabolic acidosis, and bleeding and coagulation dysfunction. Some COVID-19 patients have only mild or atypical symptoms, including, initially, even some of those who go on to develop severe and critical cases^4^ The chest computed tomography of COVID-19 patients is characterized by the ground-glass opacity and bilateral patchy shadowing^5^. For laboratory tests, it has been reported that most patients had lymphopenia and elevated C-reactive protein^6^. However, these clinical and laboratory characteristics are not easily distinguishable from pneumonia induced by infection with other common respiratory tract pathogens.

The appropriate and accurate diagnosis of the *SARS-CoV-2* infection is critical for epidemiological interventions to prevent further spread within the community^7^. Currently, molecular testing remains the only testing method for detection of virus RNA from nasopharyngeal specimens collected from suspected cases. Like many other diagnostic methods, molecular testing comes with some degree of variability with respect to sensitivity and specificity, mainly driven by the pre-analytical steps and kinetics of virus shedding from the infected individuals. For example, studies show discordant results from different types of specimens collected, naso-vs. oro-pharyngeal swabs, in COVID-19 patients^8^. Additionally, many cases that show strong epidemiologic links to *SARS-CoV-2* exposure and with typical lung radiological findings remain RNA negative in their upper respiratory tract samples. The performance of molecular tests thus depends on many factors, including the sample type^9^, the patient’s stage of infection^10^, the skill of sample collection, and the quality and consistency of the PCR assays being used. Any of these factors can lead to a substantial delay in early diagnosis and management, in turn delaying both timely life support treatment for the individual and contact tracing and preventive quarantine to contain virus spread^11^.

Antibody detection tests offer the opportunity to mitigate some of the challenges molecular testing presents. They have faster turn-around time, high throughput, and cost less per test compared to molecular testing, and thus may be a valuable adjunct where challenges to timely results and/or quality sample collection for molecular testing arise.

Serological tests for detecting *anti-SARS-CoV-2* antibodies are new to the diagnosis of Coronavirus infections. They have only rarely been utilized for diagnosis of common cold Coronavirus infections, hence many laboratories lack experience in serological testing for new *SARS-CoV-2* and may encounter initial problems with test performance characteristics and interpretation of test results in the absence of strong clinical suspicion for COVID-19.

We investigated the performance of an ELISA test for *anti-SARS-CoV-2* antibody detection, the relationship of molecular tests with serological tests in outpatient specimen and concurrently collected emergency department and pre-procedure specimens, and the dynamics of *anti-SARS-CoV-2* antibody responses in a small sample of serially-collected blood samples from inpatients with confirmed COVID-19. Further, we discuss the value and potential diagnostic and clinical use of serological test as an adjunct to molecular testing in these various cohorts.

## Methods

This study was reviewed and approved by the Baylor Scott and White Research Institute (BSWRI) Institutional Review board (IRB # 020-122)

### Study design and specimen source

This study included outpatient, emergency department, inpatient and pre-procedure adult patients from Baylor Scott & White Medical Center in Temple (Temple, TX). All adult patients were screened for symptoms of *SARS-CoV-2* infection according to WHO and Baylor Scott & White Health (BSWH) guidelines, except pre-procedure patients.

### Outpatient specimens

Serum samples from 167 patients were collected and stored at −20°C until tested. Specimens were collected from patients presenting at BSWH outpatient clinics with suspected symptoms of *SARS-CoV-2* infection. Of the 167 patients, 15 (8.9%) were confirmed positive by rRT-PCR for *SARS-CoV-2* infection, and serum specimens from rRT-PCR confirmed patients were drawn at ≥13 days after rRT-PCR test results. Additionally, 33 (19.7%) specimens were collected ≤13 days after initial rRT-PCR testing. 152 (91%) serum specimens were from patients who tested negative by rRT-PCR.

### Emergency department (ED) specimens

Nasopharyngeal and serum samples were concurrently collected from 151 patients who visited the ED with suspected symptoms of *SARS-CoV-2* infection. Specimens were collected from patients who exhibited at least one symptom related to COVID-19 disease as indicated by CDC. Symptoms included, fever, chills, cough, shortness of breath or difficulty breathing, fatigue, muscle or body aches, headache, new loss of taste or smell, sore throat, congestion or runny nose, nausea or vomiting, and diarrhea.

### Inpatient specimens

Several ED patients were transitioned to inpatient status due to clinical necessity. Residual serum specimens from 9 *SARS-CoV-2* confirmed inpatients were collected over a period of their stay in the hospital and analyzed for serum IgG.

### Pre-procedure specimens

On April 22, 2020, BSWH reopened elective surgical procedures and established a screening protocol for *SARS-CoV-2* infection. Elective surgery patients were required to submit a nasopharyngeal swab and an optional blood sample for serological testing. Accordingly, 6,271 paired nasopharyngeal swabs and blood samples were submitted by the weekend of June 12, 2020. Pre-procedure specimens can be considered truly random in distribution and represented the central Texas general population.

### Pre-COVID-19 specimens

One hundred pre-COVID-19 serum samples were selected from the BSWH specimen biobank for specificity testing of the serum IgG assay. These specimens were collected during 2018-19 Influenza season.

### Data collection and analysis

Clinical and laboratory data were extracted from electronic medical records and the laboratory information system. The receiver operating characteristic (ROC) curve plots the sensitivity against 1-specificity, or true positive rate vs. false positive rate, for all the possible cutoffs. Based on a visual assessment of the ROC curve, two potential cutoffs were chosen to calculate the sensitivity and specificity compared to rRT-PCR results. The final assay cutoff for serological testing was prepared using outpatient test results. ROC analysis and data visualization were done using EP evaluator software (Data innovations, South Burlington, VT, USA).

## Laboratory procedures

### Molecular testing for SARS-CoV-2 infection

Methods for laboratory confirmation of *SARS-CoV-2* infection were based on the rRT-PCR technique approved by the US Federal Drug and Food Administration (FDA) under an Emergency Use Authorization (EUA)^7^. Briefly, all BSWH specimens were collected either at drive through collection sites, emergency department or from inpatients using a flocked swab in Universal or Transport Media (Copan Technologies, USA). Specimens were transported at 2 – 8°C to the BSWH-Temple molecular pathology laboratory for processing and testing with less than 3 hours of transit time. The BSWH-Temple molecular pathology laboratory was responsible for *SARS-CoV-2* detection in respiratory specimens by rRT-PCR methods (Luminex Corporation, Austin, TX USA).

The *SARS-CoV-2* primers were designed by Luminex to detect RNA targets from the *SARS-CoV-2* in respiratory specimens from patients, as recommended for testing by public health authority guidelines. Luminex Aries employs primers for amplifying the *ORF1* gene and the *N* gene from the *SARS-CoV-2* virus, and the assay includes extraction and internal controls (Human RNAase P) built into the same cartridge, to verify sample lysis, nucleic acid extraction, and proper system and reagent performance. Luminex Aries offers true random-access testing, unlike the Luminex NxTAG platform, an assay for batched testing (offering high throughput capabilities) on which increased demand for testing necessitated verification and implementation. The Luminex NxTAG method also includes an additional *Envelope (E)* gene target for *SARS-CoV-2* detection.

### Antibody testing

Serum samples were collected, as stated above, from both PCR positive and negative patients, and tested for anti-*SARS-CoV-2* IgG antibodies. Testing was performed as per the instructions for use provided by the manufacturer. Briefly, the *SARS-CoV-2* IgG assay (*Ansh Laboratories,* Houston, TX, USA) uses indirect two-step immunoassay methods. In the assay, calibrators and unknowns were incubated in microtiter wells coated with purified *SARS-CoV2* recombinant antigens (spike and nucleocapsid). After incubation and washing, the wells were treated with the conjugate, composed of anti-human IgG antibodies labeled with peroxidase. After a second incubation and washing step, the wells were incubated with the substrate tetramethylbenzidine (TMB). An acidic stopping solution was then added and the degree of enzymatic turnover of the substrate is determined by wavelength absorbance measurement at 450 nm as primary test filter and 630 nm as reference filter. The absorbance measured is directly proportional to the concentration of human IgG antibodies present in the specimen. The serum IgG ELISA method was automated on Dynex DSX 4-plate instrument *(Dynex Technologies,* Chantilly, VA, USA). Calibrators and controls were run as per the manufacturer’s recommendations provided in the package insert.

The serology assay was validated and implemented as a laboratory developed test. This assay uses a three-point calibration curve. Performance characteristics were established in accordance with regulatory requirements and are available for review. During internal validations sensitivity of 95% and specificity of 98.3% were established at the time of test implementation. In this study, we used 100 pre COVID-19 specimens for additional specificity testing.

### Sample dilution experiment

In order to rule out non-specific binding, specimens that tested positive by ELISA assay were diluted using sample diluent provided in the assay kit. Specimens were diluted 1:2, 1:4, 1:8 and 1:16 and were re-tested along with an undiluted specimen. Percent recovery was calculated and plotted.

## Results

### Outpatient serological testing experience and test performance

A total of 167 serum samples were tested for serum IgG antibodies against *SARS-CoV-2;* 15 (8.9%) were positive by rRT-PCR and the remaining 152 (91%) were negative. We used these data to generate two different assay cutoffs for serum IgG and investigated the percent concordance with rRT-PCR test results.

At a lower assay cutoff, 13 AU/mL, there was a 22.3% concordance with rRT-PCR results. The percent concordance increased to 82% with increase in the assay cutoff to 35 AU/mL. Performance characteristics of the serum IgG assay were determined, including sensitivity, specificity, positive predictive value and negative predictive value, with 2 different assay cutoffs, compared to rRT-PCR (Tables 1A and 1B).

**Table 1A:**
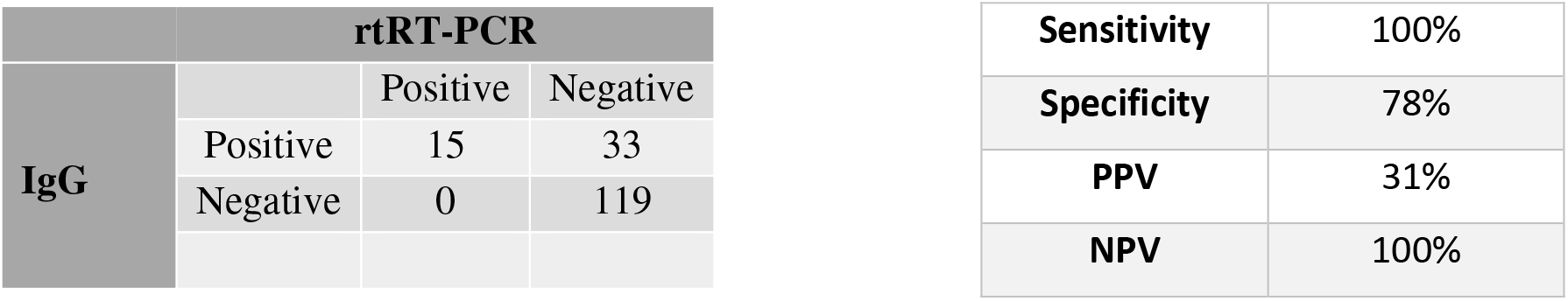
Comparison of rtRT-PCR and serum IgG using a lower (13 AU/mL) cut off

| rtRT-PCR |  |  |  |
| --- | --- | --- | --- |
| IgG |  | Positive | Negative |
|  | Positive | 15 | 33 |
|  | Negative | 0 | 119 |

|  |  |
| --- | --- |
| **Sensitivity** | 100% |
| **Specificity** | 78% |
| **PPV** | 31% |
| **NPV** | 100% |

**Table 1B:**
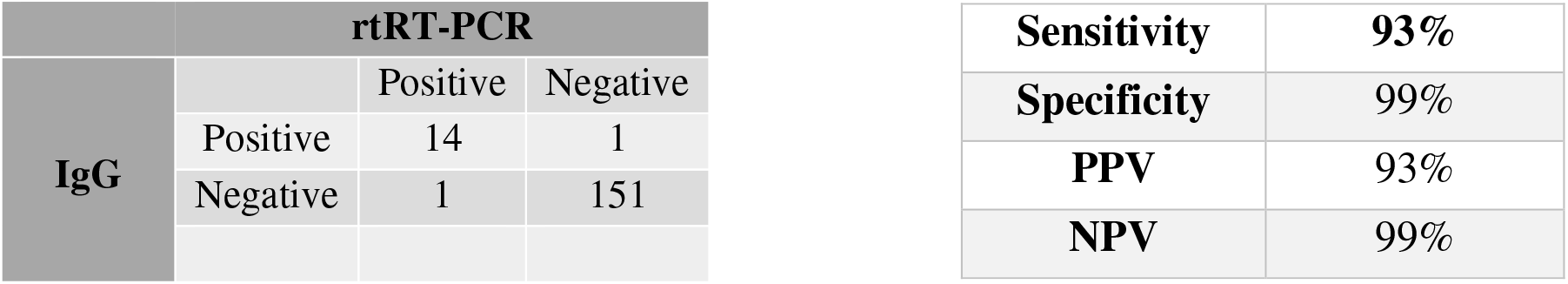
Comparison of rtRT-PCR and serum IgG using a higher (35 AU/mL) cut off Serological data collected from outpatient specimens were compared with rRT-PCR results to determine assay performance characteristics. ROC analysis was performed to draw assay cutoff, data presented Table 1A and 1B show performance characteristics at different levels of assay cutoffs.

| rtRT-PCR |  |  |  |
| --- | --- | --- | --- |
| IgG |  | Positive | Negative |
|  | Positive | 14 | 1 |
|  | Negative | 1 | 151 |

|  |  |
| --- | --- |
| **Sensitivity** | **93%** |
| **Specificity** | 99% |
| **PPV** | 93% |
| **NPV** | 99% |

Additionally, 15 serum specimens that were collected ≥13 days after initial positive rRT-PCR test had 100% concordance with the serum IgG assay, however, 33 (19.7%) serum specimens collected ≤13 days post rRT-PCR negative results were positive for *anti-SARS-CoV-2* IgG antibodies with zero percent concordance with rRT-PCR.

In order to address the discordance between rRT-PCR negative and serum IgG positive specimens and understand if the serum IgG assay had any non-specific binding issues, we retrieved five discordant serum specimens and performed a serial dilution experiment to rule out non-specific binding. Serially diluted specimens exhibited a linear decline in the antibody concentrations (Fig. 1), implying that non-specific binding was not an issue with the serum IgG assay and the discordant specimens were truly positive. From the specimen dilution experiment we were also able to determine the assay cutoff: using outpatient derived serology test results, the cutoff was set at 13 AU/mL using EP evaluator software (Fig. 2).

**Fig 1.**
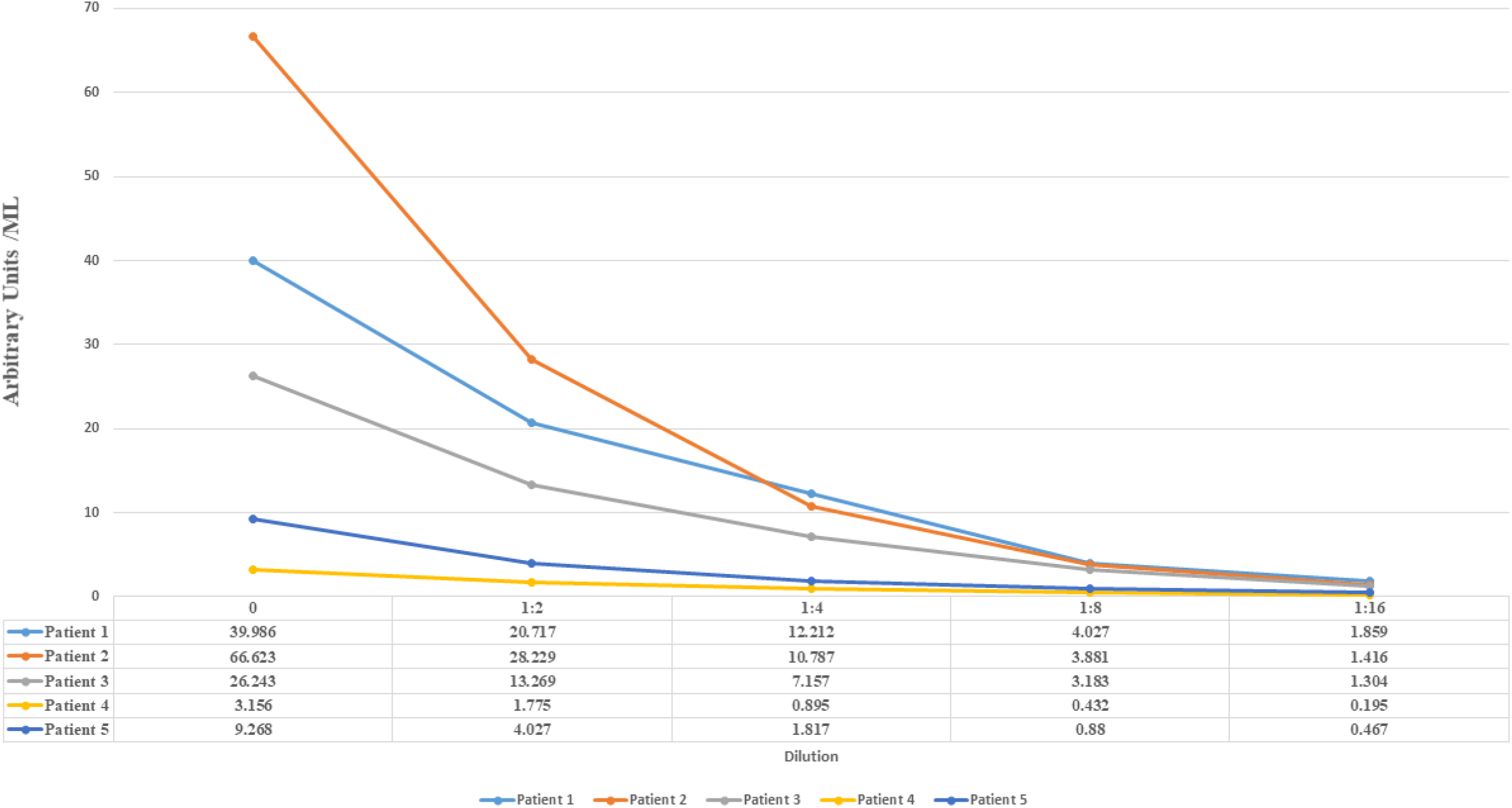
ELISA method used in the study was verified for its specificity and non-specific binding as described in the methods. Five discordant samples from outpatient specimens were serially diluted and tested on the ELISA method. Specimen tested exhibited a linear decrease in the IgG concentration and percent recovery of the analyte is shown below in the data table.

**Fig 2.**
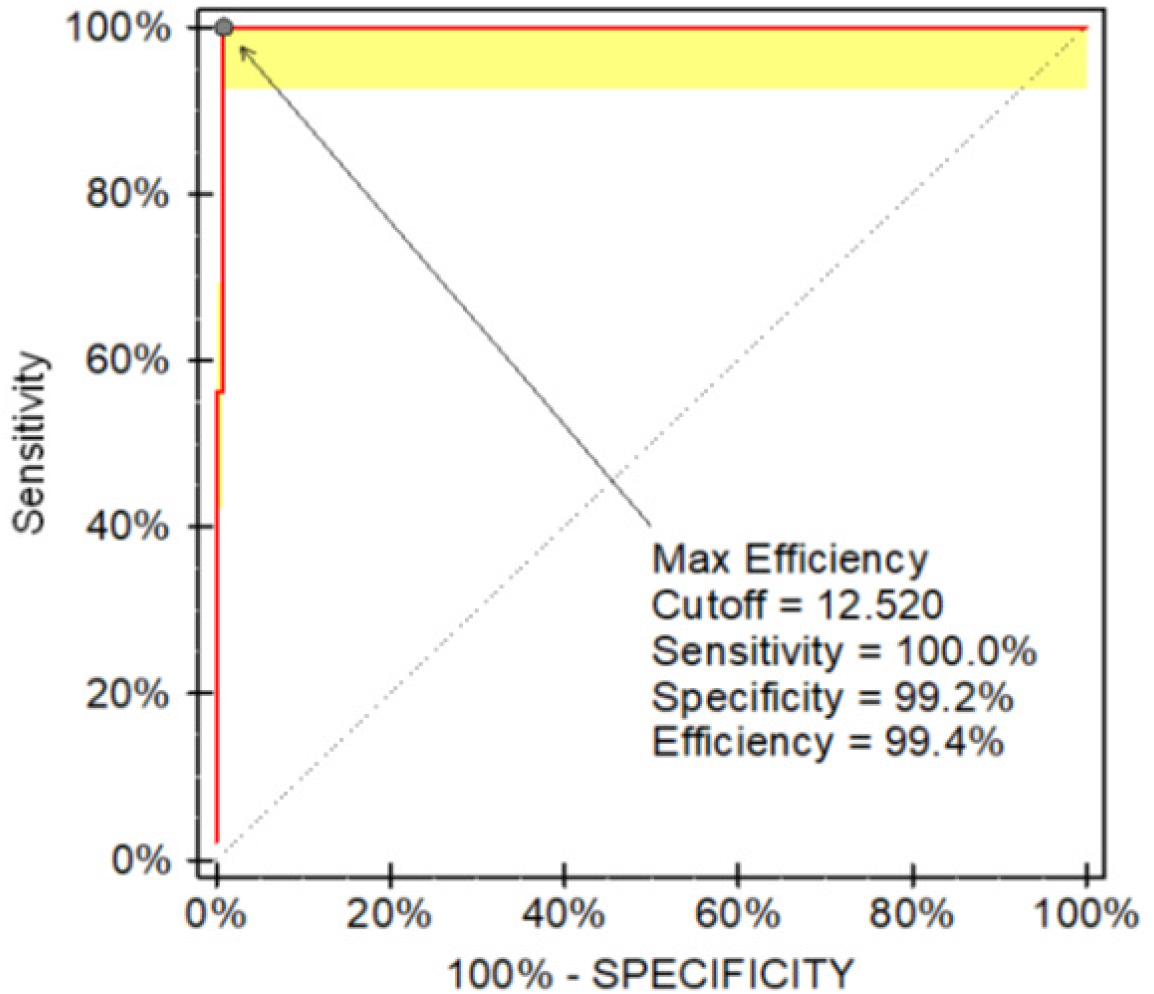
Using outpatient data and ROC analysis an assay cutoff of 13 AU/mL was calculated. ROC was determined using EP evaluator software.

Specificity of the serum IgG assay was determined using pre-COVID-19 archived serum specimens. Among the 100 pre-COVID-19 specimens, none were reactive on the IgG assay, further confirming the specificity of the assay at 100%. This also meant that negative predictive value of the assay was 100% at the lower assay cutoff.

### Emergency department serological testing experience

ED specimens, both nasopharyngeal swabs for rRT-PCR and serum for IgG assay, were concurrently collected and tested. Out of 151 nasopharyngeal swabs, 4 (2.6%) were positive by rRT-PCR and 18 (11.9%) were positive for serum IgG assay. Among the 18 patients that were positive for serum IgG, 13 (72.2%) exhibited 1-3 symptoms of COVID-19 and 5 (27.7%) patients did not present with any COVID-19 related symptoms, per CDC criteria (Table 3A). Similarly, all 4 (100%) patients that were positive by rRT-PCR had symptoms of COVID-19 disease (Table 3B). Both serology and rRT-PCR tests were negative for 53 (39.8%) and 59 (40%), respectively, patients who exhibited 1 or more COVID-19 related symptoms.

**Table 3A:**
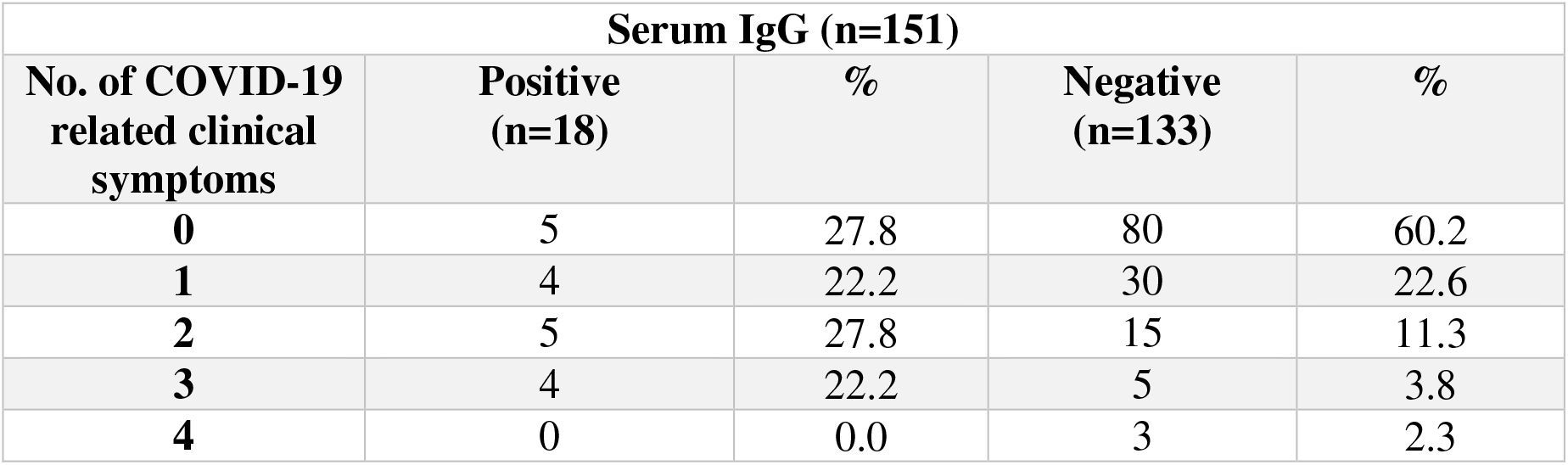
Serological and rRT-PCR testing in emergency department

**Table 3B:** Data shown in the above tables represent specimens concurrently collected from ED patients. Specimens were tested for serum IgG and *SARS-CoV-2* RNA as described in the methods. Electronic health records were reviewed for symptoms of COVID-19 as per CDC criteria and corresponding test results were noted for diagnosis of *SARS-CoV-2* infection in the ED cohort.

| rtRT-PCR (n=151) |  |  |  |  |
| --- | --- | --- | --- | --- |
| No. of COVID-19 related clinical symptoms | Positive (n=4) | % | Negative (n=147) | % |
| 0 | 0 | 0.0 | 88 | 59.9 |
| 1 | 2 | 50.0 | 33 | 22.4 |
| 2 | 1 | 25.0 | 16 | 10.9 |
| 3 | 1 | 25.0 | 7 | 4.8 |
| 4 | 0 | 0.0 | 3 | 2.0 |

Patients exhibiting COVD-19 related symptoms had a concordance of 72.2% with serum IgG assay results, providing an opportunity to consider the patients as potential *SARS-CoV-2* infections for triaging to appropriate COVID-19 designated wards if there was clinical necessity, especially in the absence of positive rRT-PCR test results and with the high negative predictive value of the serological test.

These ED data provide an opportunity for preventing spread of infection in the non-COVID-19 wards if patients were to be falsely classified as COVID-19 negative, just based on rRT-PCR results.

### Inpatient serological testing experience

Many of the ED patients were admitted as inpatients, provideing an opportunity for a limited longitudinal study on the serum IgG titers from residual specimens. Data presented in Fig. 3 show that there was a moderate to high increase in the serum IgG titers for a few patients within a very short period. Among the 9 patients that were followed, 5 demonstrated a sharp increase in the IgG titers within 1-3 days of initial testing (patient numbers 1, 2, 4, 6 and 7). There was moderate increase in the IgG titers for one patient (patient number 5) while the IgG levels had plateaued in 3 patients (patient numbers 3, 8 and 9).

**Figure 3.**
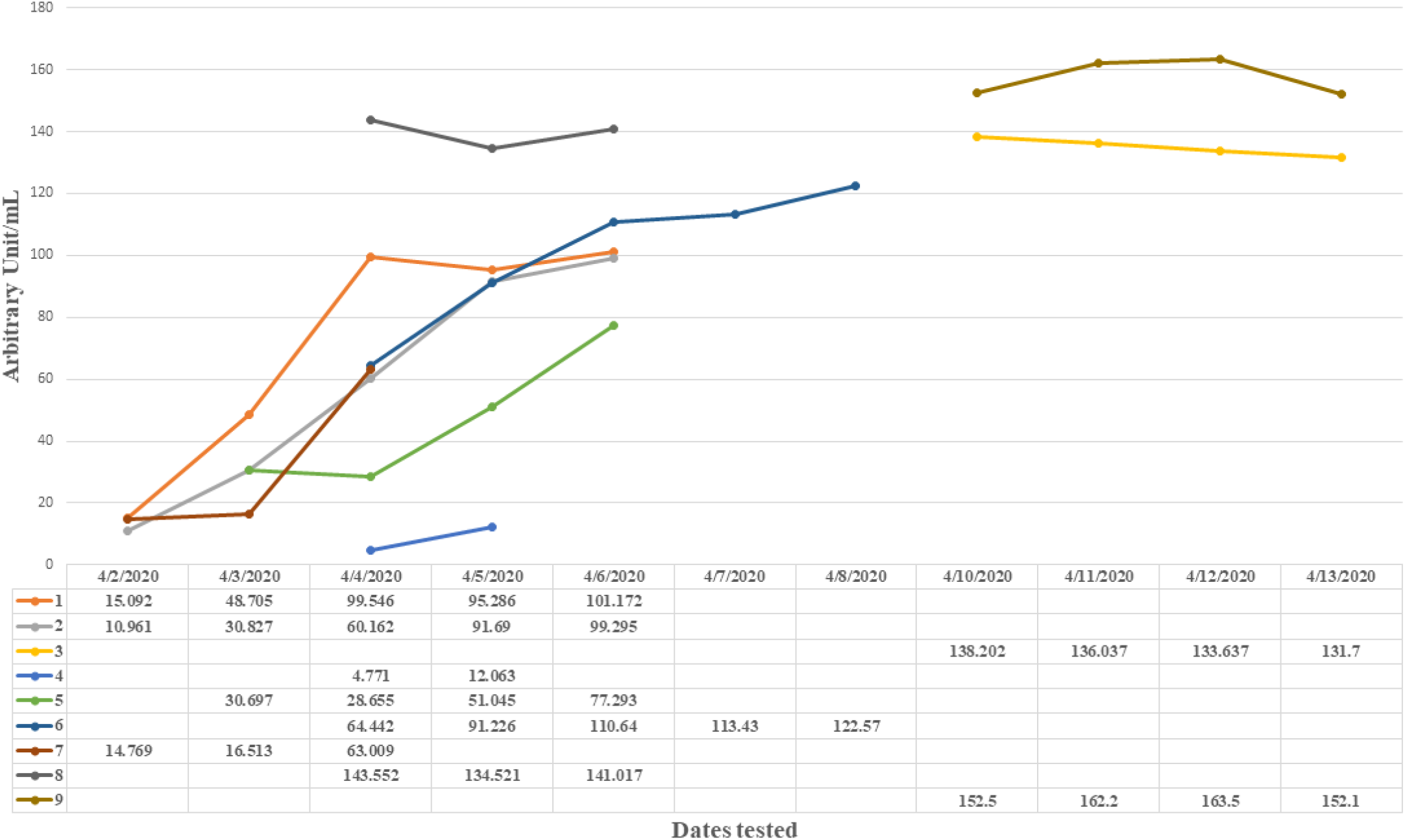
Residual serum specimens were available for 9 in-patients who were followed-up for serum IgG titers over the period of their hospital stay. Serum IgG titers were determined using ELISA method as described in the methods and AU/mL is plotted against the time. Data presented in the table represents actual titers for specific patients vs. time.

Titers of *SARS-CoV-2* antibodies can reflect the progress of viral infection. A sharp increase in the titers within a short period suggests an ongoing and active infection; therefore, these patients needed to be placed in COVID-19 designated wards to prevent cross contamination. The conventional belief, for many infections, is that a rise in the serum antibody titers corroborates an enduring infection.

### Pre-procedure serological testing experience

As mentioned in the methods, a total of 6,271 rRT-PCR and serum IgG tests were performed by the end of June 12, 2020. Among the 6,271 rRT-PCR tests, 60 (0.95%) were positive. Serum IgG test was positive for 351 (5.5%) patients among the patients who submitted paired specimens for pre-procedure screening. Since the pre-procedure patient population represented a random group from various parts of the central Texas region, we estimated seroprevalence of COVID-19 to be at 5.5% in this part of the nation.

Pre-procedure serological testing data clearly suggested a higher prevalence of COVID-19 compared to rRT-PCR data. These findings, combined with other clinical symptoms and laboratory findings, may allow careful decision making for downstream procedures, such as rescheduling or use of enhanced personal protective equipment during the invasive procedure.

### Discussion

Testing for *SARS-CoV-2* RNA has become the standard for COVID-19 diagnosis^12^. However, a number of false negative results have been reported, resulting in a failure to quarantine infected patients^12^. If unchecked, this could cause a major setback in containing viral transmission^13^. Serological tests are crucial tools for assessments of *SARS-CoV-2* exposure, infection and potential immunity. Their appropriate use and interpretation requires accurate assay performance data^14^

We described the use of serological testing for *SARS-CoV-2* infection in various healthcare contexts, examining outpatient, emergency department, inpatient, and pre-procedure patients. Outpatient data were utilized to determine the performance characteristics of the IgG ELISA assay, which was validated and implemented as a laboratory-developed test. Two assay cutoffs were established based on its performance compared to rRT-PCR method. Both the cutoffs provided a nearly 100% negative predictive value.

As shown by many studies, rRT-PCR results have been variable due to pre-test probabilities, however, serological tests with high specificity and negative predictive value can be used as an adjunct to rRT-PCR findings, combined with other clinical symptoms and laboratory findings for appropriate patient care.

Increased virus shedding and transmission have been reported in people asymptomatic for COVID-19^10^. rRT-PCR findings combined with serological testing can further delineate different types of infected people, including asymptomatic individuals. We have shown here that both pre-procedure (asymptomatic) and ED (symptomatic) patients had higher positivity rates by serological testing than rRT-PCR, including several of ED patients who were symptomatic as per the CDC definition of COVID-19.

Interestingly, it has been shown that the typical incubation period for *SARS-CoV-2* infection could be anywhere between 2 and 14 days, with the average being 7 days^3,8^ By the time patients show serious signs and symptoms, like shortness of breath, it could be greater than 7 days post-infection when these patients present to ED, a point at which virus load in nasopharyngeal swabs could be below detectable levels. The long incubation time allows for antibody development in the infected individual, and both serum IgM and IgG would begin to appear at levels detectable by commercial assays^13^. It is therefore important to have an assay that has high specificity, sensitivity and negative predictive value, such as the one used here. In this study, 13 (72%) out of 18 of the patients in ED were found to have COVID-19 specific symptoms and tested positive for serum IgG, indicating an ongoing infection.

Our inpatient data were indicative of current or ongoing infection with *SARS-CoV-2,* based on the increase in the titer of specific antibodies. Several infectious diseases are diagnosed based on serological tests and the diagnosis is often based on the demonstration of an increase in the serum antibody levels between acute and convalescent specimens^15^. We strongly believe that a similar diagnostic approach is necessary for a largely unknown entity such as COVID-19, perhaps with a short duration between specimens collected, as demonstrated in this study.

High specificity testing is crucial in low-prevalence settings, as shown in our data; the ELISA test we employed had 100% specificity with a negative predictive value of 100%. We evaluated 6000+ serum and nasopharyngeal swabs from pre-procedure patients and found that the positivity rate was significantly higher by serological test than rRT-PCR. These preprocedure patients were asymptomatic and represented a true random sample from the central Texas region. We observed 5.5% sero-prevalence in this region. Serological tests thus have a significant role in downstream clinical decision-making (use of enhanced PPE or rescheduling) and patient triaging to appropriate care and/or discharge.

The intent of this study was not to provide any guidelines or recommendations on how to use anti-*SARS-CoV-2* serological tests in various settings, especially since the CDC recommends that serological tests alone should not be used for diagnosis. However, the CDC also recommends that, in certain situations, serologic assays may be used, in conjunction with viral detection tests, to support clinical assessment of persons who present late in their illnesses^3^. We recommend that it is best left to the discretion of individual healthcare facilities and the preference of scientific community as to what the specific downstream applications of serological tests may be in the management of COVID-19.

### Conclusion

We demonstrated how an adjunct serological test with high negative predictive value for *SARS-CoV-2* infection can be leveraged for appropriate clinical decision making in various clinical scenarios.

## Acknowledgment

Authors sincerely thank Jeffry Hunt for help with data extraction, Courtney Shaver for statistical help and Briget Da Graca for editorial help.

